# Phonic Tics in a Rat Model of Tourette Syndrome Enable Research on Symptom-Based DBS

**DOI:** 10.1101/2023.08.06.551271

**Authors:** Boriss Sagalajev, Lina Lennartz, Niloofar Mokhtari, Mikolaj Szpak, Thomas Schüller, Juan Carlos Baldermann, Pablo Andrade, Veerle Visser-Vandewalle, Thibaut Sesia

## Abstract

The lack of a rodent model for both motor and phonic tics hinders research on deep brain stimulation (DBS) for refractory Tourette syndrome (TS). Striatal disinhibition with a GABA-A antagonist (bicuculline) was previously shown to induce tic-like hyperkinesia in rats and monkeys, while tic-like vocalizations were studied and confirmed only in the latter. We, therefore, assessed whether vocalizations accompany hyperkinesia also in rats and whether they respond to thalamic DBS. Rats underwent surgical implantation of a unilateral guide cannula targeting the caudate putamen (CPu) or nucleus accumbens (NAc). Additionally, they were implanted with an ipsilateral stimulation electrode targeting the border between the central medial (CM) and ventrolateral (VL) thalamic nuclei. Motor changes and ultrasound vocalizations were recorded and characterized offline. Bicuculline in CPu led to the development of hyperkinesia in the form of arrhythmic, myocloniform shoulder jerks sporadically alternating with episodes of dystonic lordosis. Likewise, bicuculline in NAc resulted in hyperkinesia, but at a much smaller dose and in conjunction with nonsensical vocalizations. DBS of CM/VL, but not adjacent regions, attenuated hyperkinesia. Also, early results indicate that thalamic DBS attenuates vocalizations, yet the hotspot for stimulation remains to be determined. In conclusion, striatal disinhibition leads not only to the development of hyperkinesia but also vocalizations in rats. The resemblance of hyperkinesia to motor tics and vocalizations to phonic tics is evident in their appearance and susceptibility to DBS. Rats can, therefore, be used to study DBS for symptom-based TS therapy.

## Introduction

Tourette syndrome (TS) is a debilitating disorder characterized by unwanted sudden, repetitive, arrhythmic movements and vocalizations. They are commonly referred to as tics and differ from other involuntary movements in that they are often preceded by a premonitory urge and followed by a sense of relief [1]. Although patients can suppress tics, this process is exhausting and can only be sustained for a limited amount of time. TS affects 8 per 1000 children [2] and is often accompanied by neuropsychiatric comorbidities such as attention deficit hyperactivity disorder and obsessive-compulsive disorder. Additionally, mood disorders such as anxiety and depression are common among TS patients [3]. Current treatment consists of a comprehensive behavioral intervention for tics [4] and of pharmacological therapy with antipsychotics, noradrenergic agents, and cannabinoids, among others [5]. While TS symptoms usually first appear in early childhood and dissipate by late adolescence, some patients continue to be severely affected even after reaching adulthood and receiving treatment. In the most severe cases, these patients resort to methods of functional neurosurgery, such as deep brain stimulation (DBS) [6]. Although DBS has proven its efficacy in movement disorders like Parkinson disease through large, randomized trials [7], evidence for its efficacy in treating tics remains limited due to the low prevalence of patients with refractory TS, with most publications in the field being case reports [8]. Also, DBS mechanisms for TS remain largely unknown [9, 10]. Therefore, the establishment of an animal model with the pathophysiology, behavioral phenotype, and treatment responsiveness like that of TS patients would represent a major step forward in elucidating DBS underpinnings with methods unsuitable for clinical application [11]. In DBS for Parkinson, for example, models relying on pharmacological disruption of the nigrostriatal pathway led to a thorough understanding of DBS mechanisms and further refinement of its protocols [12].

Striatal GABAergic interneurons integrate input from a wide cortical area and exert selective inhibition on striatal projection neurons [13]. They, therefore, act as an amplifier of signal-to-noise ratio between the cortex and striatal targets. Disruption of their function may, therefore, create a condition where an increase in cortical activity may induce hyperkinesia, in such forms as chorea [14], dyskinesia [15], myoclonia [16], convulsions [17], and dystonia [18]. More than a decade ago, disinhibition of the striatum with a GABA-A antagonist (bicuculline) was first suggested as a TS animal model [19]. However, while tic-like movements and vocalizations have since been observed in monkeys [20], only the former have been studied and verified in rats [21-24]. We, therefore, set out to determine if rats develop tic-like vocalizations too. Also, we sought to examine if tic-like behavior responds to DBS of the central medial (CM) and ventrolateral (VL) thalamic nuclei [25]. We argue that this region in rats represents the closest equivalent to one of the most common DBS targets in TS patients [8]: the area formed by the thalamic centromedian nucleus and nucleus ventrooralis internus [26, 27].

## Materials and methods

### Ethical approval

All experiments were approved by the local regulatory authority (*Landesamt für Natur, Umwelt und Verbraucherschutz Nordrhein-Westfalen* (LANUV), North Rhine-Westphalia, Germany; permission no. 81-02.04.2019.A066) and were carried out in accordance with the EU Directive 2010/63/EU on the protection of animals used for scientific purposes.

### Animals

Sprague Dawley male rats aged 2-3 months were obtained from Janvier Labs (Le Genest-Saint-Isle, France). They were kept together in cages littered with low-dust spruce granulate and enriched with cardboard tubes, paper towels, and aspen bricks. Access to food and water was ad libitum. The cages were held in an animal facility with a normal day-night cycle (12:12 hours).

### Stereotactic surgery

Thirty minutes before surgery, rats received subcutaneous injections of two analgesics: carprofen (5 mg/kg) and buprenorphine (0.05 mg/kg). Isoflurane anesthesia was then induced at 4% and maintained at 2% (1 l/min). Steady breathing, pink skin coloration, and absence of interdigital reflex ensured an appropriate anesthesia level. The rat’s temperature was monitored with a rectal probe and maintained with a heating pad. The head was secured to a stereotactic frame and the eyes were moistened with a lubricant. The scalp was shaved, disinfected with iodine, and anesthetized with subcutaneous 2% lidocaine (B. Braun, Melsungen, Germany). After cutting the skin and removing underlying muscles, a hole was drilled in the skull to insert a guide cannula (PlasticsOne, Roanoke, Virginia, USA). To target the dorsal striatum (caudate putamen, CPu), we positioned the cannula’s tip 1.5 mm rostrally from the bregma, 2.5 mm laterally from the midline, and 3.0 mm deep from the dura. In a different group of rats, to target the ventral striatum (nucleus accumbens, NAc), we positioned the cannula’s tip 1.8 mm rostrally from the bregma, 1.8 mm laterally from the midline, and 5.7 mm deep from the dura [25]. An additional hole was drilled for the implantation of a concentric electrode (PlasticsOne). To target the CM/VL border, we positioned the electrode’s tip 2.4 mm caudally from the bregma, 0.9 mm laterally from the midline, and 6.4 mm deep from the dura [25]. To mimic clinical practice, the electrode was implanted along the longest VL axis [28]. All rats had a maximum of two implants: one cannula plus one ipsilateral electrode. Half of the rats had implants placed in the left hemisphere and the other half in the right hemisphere. The implants were anchored to the skull with micro-screws and coated with dental cement. Then the skin was stitched, and the rat was returned to its cage. The guide cannula was protected with a dummy cannula and the electrode with a dust cap. Behavioral testing did not start until full recovery (10 days after surgery). After it, the brains were collected for Nissl staining. Lesions from the implants [29, 30] on coronal slices indicated whether they were implanted correctly.

### Brain injections and stimulation

Rats were split into two groups depending on the injection site: CPu or NAc. Each group was then further split into two subgroups depending on whether the rats received bicuculline (Merck, Darmstadt, Germany) or vehicle (artificial cerebrospinal fluid, aCSF). Each rat underwent three DBS sessions at three different intensities (0, 100, or 200 μA) in random order at least two days apart. Each time, the DBS session was preceded by a striatal injection.

The injections were done using a gas tight syringe (Hamilton, Bonaduz, Switzerland) connected through a plastic tube (PlasticsOne) to an internal cannula (PlasticsOne). The latter, in turn, was temporarily inserted into the chronically implanted guide cannula and protruded 1 mm from its tip. The drug was then infused with a syringe pump at a volume of 1 μl over a period of 2 min (Pump 11 Elite, Harvard Apparatus, Holliston, Massachusetts, USA). After the infusion, the internal cannula remained inside the guide cannula for an additional 30 s to prevent backflow. The estimated extent of diffusion from the internal cannula’s tip was between 0.8 and 1.1 mm [31].

DBS was applied using a current source (STG 4004, Multi Channel Systems, Reutlingen, Germany) connected through a flexible cable (PlasticsOne) to the chronically implanted concentric electrode. Biphasic pulses (−100/+100 μs) were delivered continuously at 100 Hz. An intensity of 0 μA was used for sham DBS, while 100 and 200 μA for real DBS. The estimated range of brain excitation from the electrode’s tip was 0.5 mm at 100 μA and 0.75 mm at 200 μA [32, 33]. We previously used similar DBS parameters for improving spatial memory in a rat model of Alzheimer disease [30].

### Behavioral testing

Immediately after striatal injection, the rat was placed into a chamber with four grey walls and a black floor. The chamber’s width, length, and height were 60 cm. Using a near-infrared camera (Basler AG, Ahrensburg, Germany) two meters above the chamber, the rat was observed on a computer. The computer and other surroundings were hidden from the rat behind a grey curtain. Additionally, an ultrasound microphone (Noldus, Wageningen, Netherlands) was used to record rat vocalizations. During the test, the rat was let free to explore the chamber. All rats were habituated to experimental conditions for at least two days.

### Data analysis

We used EthoVision XT 15 software (Noldus) to manually mark the onset and offset of every hyperkinetic movement in recorded videos. Ultrasound vocalizations were detected in a spectrogram created by passing the audio signal through the Fourier transform in UltraVox XT 3.2 (Noldus). Graphs were made in Python (version 3.9.13) using Seaborn (version 0.12.2) and Matplotlib (version 3.7.0) libraries. Statistical analysis was done in Python using Pingouin (version 0.5.3) and SciPy (version 1.9.1) libraries.

For the statistical analysis of the DBS effect, the data were normalized, when necessary, using the square root transformation to fulfill the assumptions of normality and equal variance. DBS-induced changes were then compared using a one-or two-way ANOVA. Results were considered significant at a p-value below 0.05 when statistical power (1–β) exceeded 0.8 at a significance level (α) of 0.05 [34].

## Results

### Effective dose of bicuculline

To prevent unwanted side effects, bicuculline was administered at a dose that would induce motor or phonic tic-like behavior in 75% of the rats (ED75). ED75 in CPu was determined to be 500 ng, with an effect duration of 80 min. DBS was, therefore, applied 10 min after the injection for 60 min. In contrast, ED75 in NAc was determined to be 25 ng, with an effect duration of 15 min. Consequently, DBS was applied 5 min after the injection for 5 min.

### Hyperkinesia during sham DBS

While aCSF in CPu and NAc did not affect rat behavior, bicuculline led to the immediate development of hyperkinesia in the form of myoclonic shoulder jerks, which occasionally alternated with episodes of dystonic lordosis during sham DBS (**Video S1**). While myoclonic movements lasted up to one second, duration of dystonic movements ranged from just over a second to almost a minute. Interestingly, myoclonia and dystonia forced the upper body to rotate in the contralateral direction to the injection side (left vs right striatum). Yet, the rats looked and walked predominantly in the opposite direction, suggesting an adaptation response to maintain their gait and posture.

A stacked histogram showing how long hyperkinesia lasted every 20 seconds (**Fig. 1A**) revealed that the mean duration of hyperkinesia fluctuated between 2 and 6 seconds depending on the post-injection time of bicuculline and was solely characterized by myoclonia. However, whenever the duration of hyperkinesia exceeded two standard deviations from the mean, the rats developed dystonia. Thus, our data indicate that myoclonia and dystonia did not represent two independent forms of hyperkinesia but rather different levels of severity.

**Figure 1.**
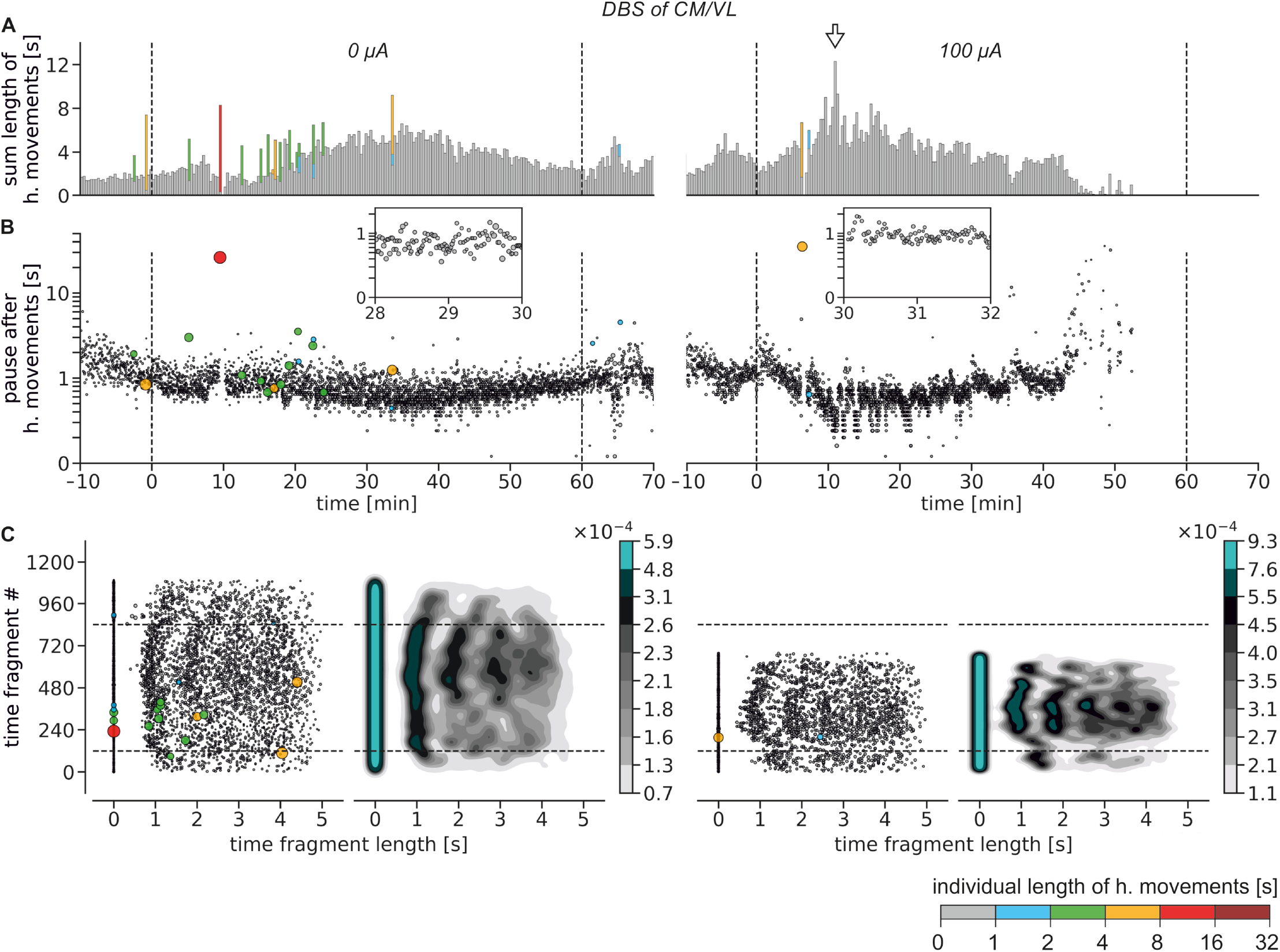
Hyperkinesia from CPu disinhibition. Example from a single rat. **A:** Stacked histograms depicting the sum length of all hyperkinetic (h.) movements every 20 s (bin width). The arrow indicates a moment when the rat did not develop dystonia during real DBS, unlike during sham DBS, when the duration of hyperkinesia exceeded two standard deviations from the mean. **B:** Bubble plots depicting the occurrence pattern of hyperkinetic movements as a function of the post-movement pause. The insets show a zoomed-in period of myoclonia forming fading bursts. **C:** Peri-event rasters of experimental sessions shown in panel B with respective heatmaps created using kernel density estimation. Vertical dashed lines in panels A and B, as well as horizontal dashed lines in panel C, denote the onset and offset of DBS. Bicuculline was injected 10 min before DBS. Gray bars and bubbles represent myoclonia, while colored bars and bubbles represent dystonia.

A bubble chart depicting how long the pause after each hyperkinetic movement lasted (**Fig. 1B**) revealed that immediately after bicuculline injection the occurrence rate of myoclonia was steady. However, as bicuculline took full effect, their rate started to drop and then rise again. Myoclonia, thus, formed episodes of fading bursts indicating the development of system fatigue (neuronal or muscular). Dystonia, in contrast, occurred sporadically, yet it occasionally heralded a temporary cessation of hyperkinesia, further supporting the notion of growing fatigue.

To assess whether hyperkinetic movements followed a certain rhythm, we drew a peri-event raster depicting the probability of their occurrence within the first 5 s after each other. Also, for better visualization of movement clusters, a heatmap of the raster was created using kernel density estimation (**Fig. 1C**). The two plots revealed that hyperkinetic movements occurred on average every second, yet the deviation from the average grew with each successive movement. Thus, rather than following the activity of a possible internal pacemaker, the exact timing of the hyperkinetic movement depended largely on when it occurred beforehand.

### Hyperkinesia during real DBS

Real DBS of CM/VL was applied to attenuate hyperkinesia resulting from CPu disinhibition. In contrast to sham DBS (see above), hyperkinesia during real DBS was characterized by myoclonia that appeared even when duration of hyperkinesia exceeded two standard deviations from the mean, suggesting that they resulted in such instances from attenuation (i.e., fragmentation) of dystonia (**Fig. 1A**). Associated fatigue, however, remained largely unaffected since myoclonia still formed episodes of fading bursts and sporadic dystonia still heralded a temporary cessation of hyperkinesia (**Fig. 1B**). Also, arrhythmicity of hyperkinetic movements remained unchanged, indicating that DBS did not entrain them (**Fig. 1C**).

For statistical analysis of the DBS effect, we compared the change in the summed length of all hyperkinetic movements. However, because bicuculline’s effect in CPu lasted for 80 min, we focused on analyzing 5-min intervals taken immediately before and after DBS onset and offset, as well as those taken 20 and 40 min into DBS. This was done to expedite the analysis process. Thus, a one-way repeated measures ANOVA showed that DBS of CM/VL attenuated hyperkinesia at the end of its 60-min application (**Fig. 2**, F_2,14_=11.110, p<0.001, 1–β=0.965, α=0.05, n=8). Bonferroni’s post-hoc test against the sham intensity (0 μA) showed that the DBS effect was significant at 100 μA (p<0.001) but not at 200 μA (p=0.210). Notably, the DBS effect persisted after its offset (F_2,14_=9.732, p=0.002, 1–β=0.935, α=0.05, n=8), with the effective intensity again being 100 μA (p=0.001) but not 200 μA (p=0.062).

**Figure 2.**
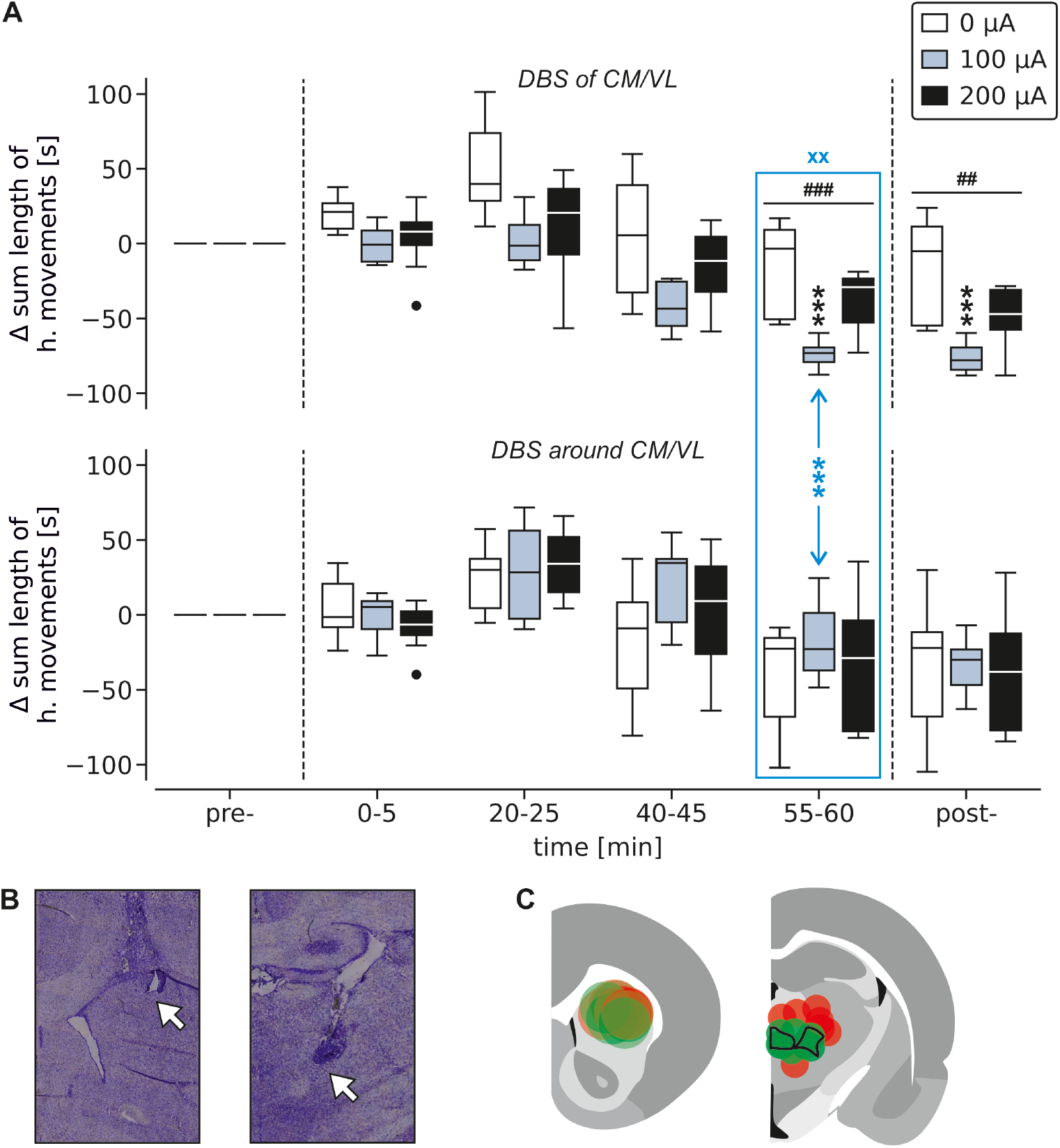
DBS effect on CPu hyperkinesia. **A:** DBS of CM/VL, but not of the adjacent regions, attenuated hyperkinesia (^###^ p<0.001, ^##^ p<0.01, one-way repeated measures ANOVA; ^xx^ p<0.01, two-way mixed ANOVA, interaction; *** p≤0.001, Bonferroni’s post-hoc test). Boxes – interquartile range, lines – median, whiskers – min and max values, circles – outliers. **B:** Implantation sites of the guide cannula (left) and DBS electrode (right) were verified in coronal brain slices after Nissl staining. Arrows show where the tip of the implants was. **C:** Estimated spread of bicuculline injections at 1 μl (left) and DBS field at 100 μA (right) [31-33]. While all rats received bicuculline in CPu, half of them (n=8, green) underwent DBS of CM/VL (outlined in black) and the other half (n=7, red) underwent DBS of the adjacent regions.

In the CPu group, DBS was also applied in regions adjacent to CM/VL, including the posterior thalamus (n=4), mediodorsal thalamus (n=2), and dorsal hypothalamus (n=1). When these rats were considered together, DBS failed to attenuate hyperkinesia across all time intervals (p>0.2, one-way repeated measures ANOVA). Yet, the fact that 1–β consistently remained below 0.8 with an α of 0.05 suggests that the analysis might not have had enough power to detect a significant effect [34].

To assess if DBS of CM/VL was more efficacious than DBS of the adjacent regions, we conducted a two-way mixed ANOVA for the last 5-min interval of the DBS duration. It showed a significant interaction between the DBS region and intensity (F_2,26_=7.851, p=0.002, 1–β=0.899, α=0.05, n_CM/VL_=8, n_adjacent regions_=7). Bonferroni’s post-hoc test showed that the difference between brain regions at 0 μA remained insignificant (p=0.102), indicating that during this time interval bicuculline effect was equally strong in both groups. Also, it showed that the difference between brain regions was significant at 100 μA (p<0.001) but not at 200 μA (p=0.822).

Bicuculline in NAc did not always lead to the development of hyperkinesia. Instead, in some cases, it only led to the development of vocalizations (**Fig. 3**). Consequently, a one-way ANOVA remained underpowered due to the small sample size (n_0 μA_=4, n_100 μA_=4, n_200 μA_=3), preventing detection of significant changes in the summed length of hyperkinetic movements during and after DBS of CM/VL (p>0.9, 1–β=0.05, α=0.05).

**Figure 3.**
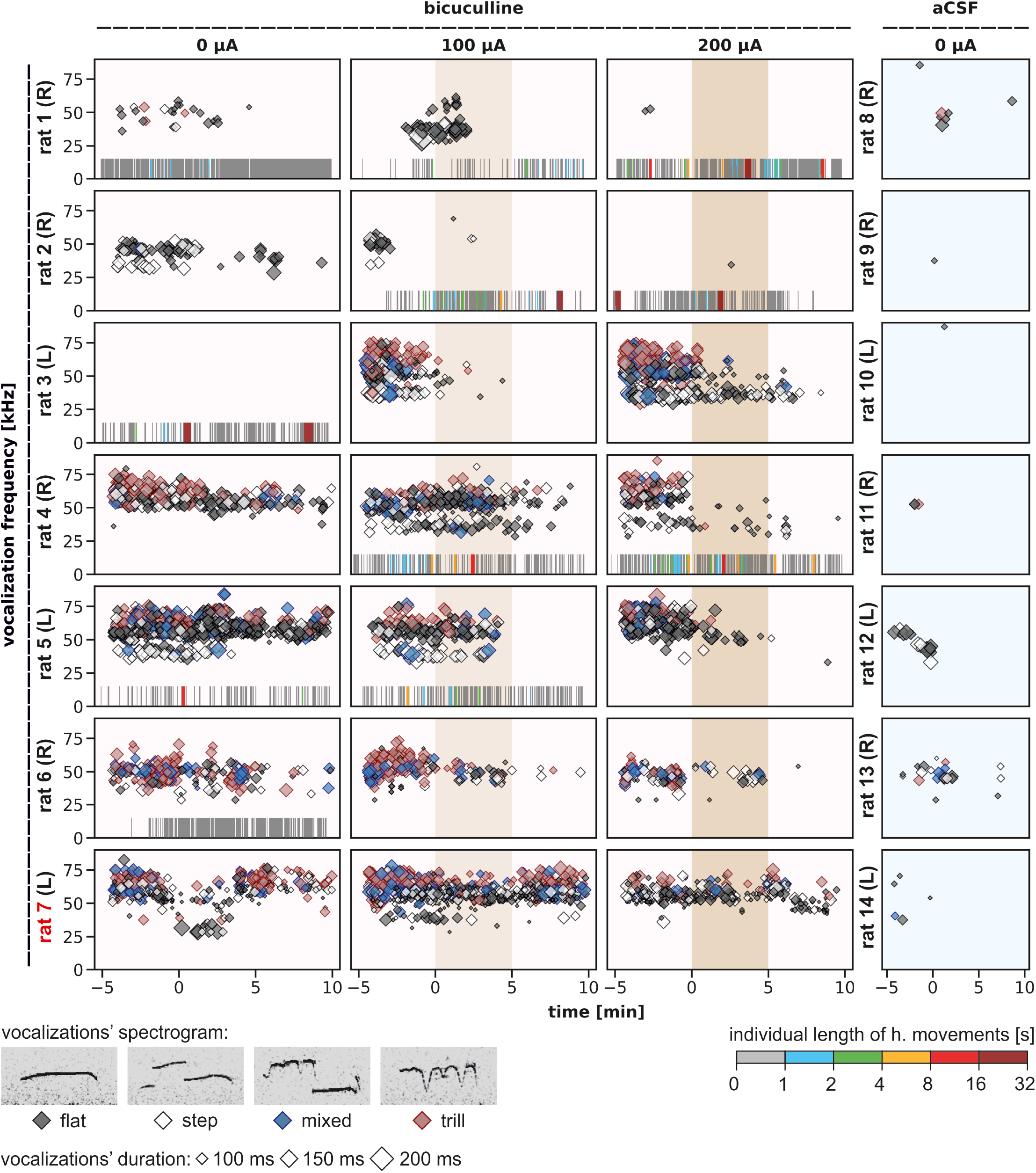
Vocalizations from NAc disinhibition. Colored diamonds depict vocalizations of different classes based on their shape in the spectrogram: flat, step, mixed, or trill. Vertical stripes at the bottom of individual plots depict instances when vocalizations were accompanied or replaced by hyperkinetic (h.) movements. Gray stripes represent myoclonia, while colored stripes represent dystonia. All rats had only one cannula and one ipsilateral electrode in the left (L) or right (R) hemisphere. Rats 1-6 underwent DBS of CM/VL, while rat 7 underwent DBS of an adjacent region (see **Fig. 4** for implant localization). DBS was applied from 0 to 5 min. Injections were performed 5 min before DBS.

Normal motor behavior in rats that received aCSF in CPu (n=6) or NAc (n=6) remained unaffected by DBS of CM/VL at all tested intensities (0, 100, 200 μA).

### Vocalizations during sham DBS

During the first 20 min after the rats received bicuculline in CPu, they remained practically silent. Ultrasound recordings, therefore, stopped, while video recordings continued for another 60 min until the end of hyperkinesia. In contrast, after the rats received bicuculline in NAc, they began vocalizing immediately. Like hyperkinesia, vocalizations lasted for about 15 min during sham DBS (**Video S1**); however, they did not always develop together. With one injection per day, the same rat could develop both or only one of them (**Fig. 3**).

Rat vocalizations were classified into two categories based on their shape in the spectrogram: flat or frequency-modulated calls. Additionally, the latter category was further divided into step, mixed, or trill calls, as previously described [35]. While flat calls are emitted during social interactions, frequency-modulated calls are emitted in affective states. In our study, however, rats that received aCSF in NAc remained mostly silent, while rats that received bicuculline in NAc developed vocalizations of all four shapes. Apart from the shape, the audio frequency of vocalizations can also indicate the rat’s mood, with vocalizations at 18-32 kHz denoting negative emotions and those at 35-80 kHz denoting positive emotions [36]. We found that vocalizations from the same bicuculline injection in the same rat could rapidly alternate between these two frequency ranges. Thus, both the shape and frequency of vocalizations suggest that they were nonsensical.

### Vocalizations during real DBS

Real DBS of CM/VL was applied to attenuate vocalizations resulting from NAc disinhibition (**Fig. 4**). Out of the seven rats that received bicuculline, the electrode was implanted on target (CM/VL) in six rats, while in one rat, it was implanted off target (posterior thalamus). The latter rat was, therefore, excluded from the analysis. All four shapes of vocalizations were analyzed together, as there appeared to be no change in their ratio when DBS was applied. A two-way ANOVA was used to compare changes in the normalized (√) summed length of vocalizations during sham vs real DBS. It showed that the main effects of time (F_1,24_=4.894, p=0.037, 1–β=0.463, α=0.05) and intensity (F_2,24_=4.100, p=0.029, 1−β=0.536, α=0.05) were significant, while the interaction between them was not (F_2,24_=0.161, p=0.852, 1–β=0.050, α=0.05). Bonferroni’s post-hoc test against the sham intensity (0 μA) showed that the DBS effect was significant at 200 μA (p=0.017) but not at 100 μA (p=0.278). However, because 1–β for the two factors and their interaction remained below 0.8, the results should be considered preliminary [34].

**Figure 4.**
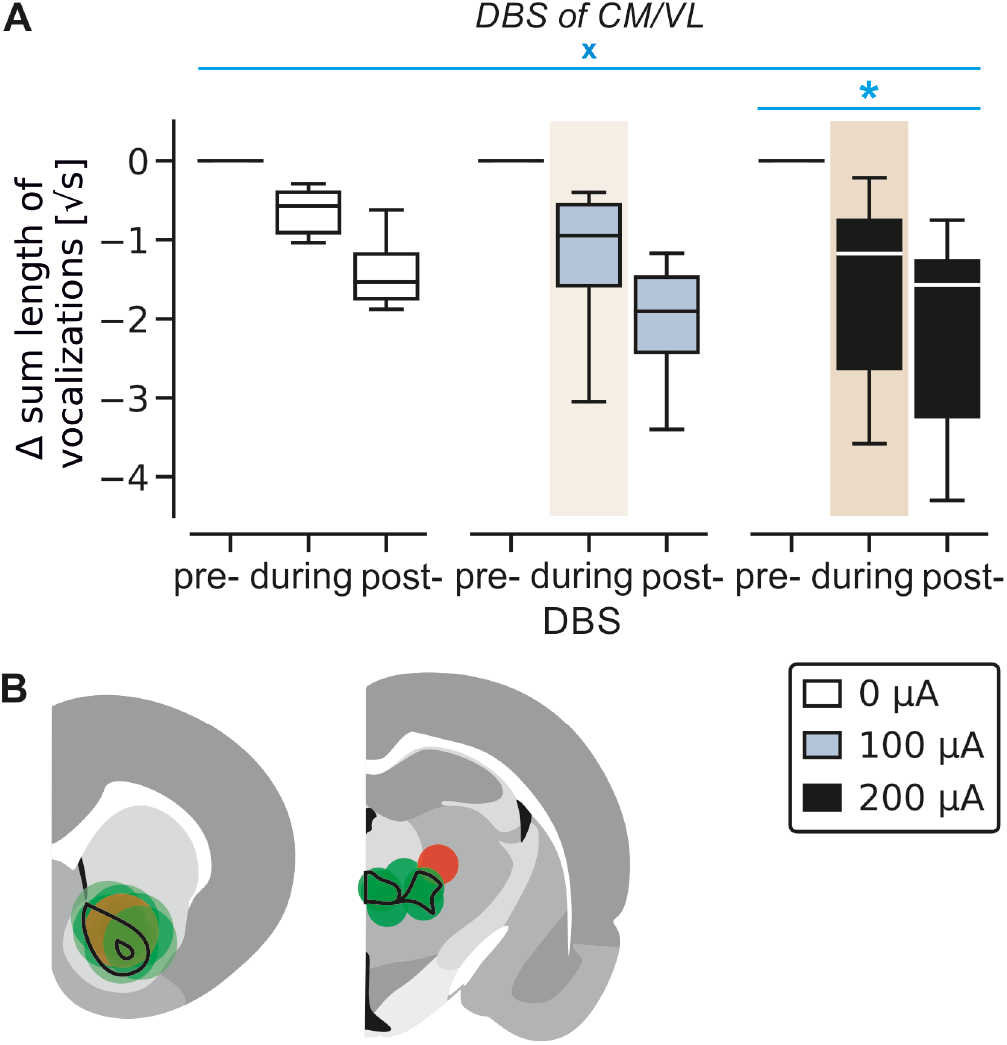
DBS effect on NAc vocalizations. **A:** Attenuation of vocalizations by DBS of CM/VL (^x^ p<0.05, two-way ANOVA, main effect of intensity; * p<0.05, Bonferroni’s post-hoc test, 0 vs 200 μA). Boxes – interquartile range, lines – median, whiskers – min and max values. **B:** Estimated spread of bicuculline injections at 1 μl (left) and DBS field at 100 μA (right) [31-33]. The green color represents rats (n=6) that underwent DBS of CM/VL (outlined in black), while the red color represents a single rat that underwent DBS of an adjacent region (excluded from the analysis). All rats had bicuculline in NAc (outlined in black).

## Discussion

Our study confirms that bicuculline-induced disinhibition of CPu and NAc leads to the development of hyperkinesia in the form of myoclonic shoulder jerks and dystonic lordosis [21-23]. Additionally, we show that disinhibition of NAc, but not CPu, leads to the development of vocalizations in rats, which has previously been shown only in monkeys [20]. Bicuculline ED75 was several times smaller in NAc than in CPu, suggesting that the former represents the striatal center for the generation of tic-like behavior. However, given the brief duration of the ED75 action in NAc, chronic bicuculline injections may be more suitable for modeling a tic disorder, as previously suggested [22]. While CPu injections always resulted in hyperkinesia, NAc injections resulted in hyperkinesia that could be accompanied or replaced by vocalizations. We, therefore, think that the exact repertoire of tic-like behaviors in the NAc group depends largely on how bicuculline spreads during the injection. Spreading in the dorsal direction (i.e., towards CPu) likely leads to hyperkinesia, in the ventral direction to vocalizations, and in both directions to their combination. The notion that NAc plays a major role in tic generation is supported by the fact that it represents one of the brain’s dopaminergic centers and that blockers of dopamine D2-receptors are used clinically for tic attenuation [5]. Also, DBS of NAc is reported to attenuate tics [37-41], although not in all patients [42].

For involuntary behavior to be classified as a tic, it must be arrhythmic, suppressible, preceded by a premonitory urge, and followed by a sense of relief [1]. In our rats, hyperkinesia was indeed arrhythmic. It took the form of myoclonia and dystonia similarly to how motor tics develop in TS patients [43]. A recent study in monkeys, however, has demonstrated that striatal bicuculline, aside from myoclonia, may also lead to the development of focal and generalized seizures, depending on the injection volume [44]. Thus, we cannot exclude that dystonia in our rats represented, in some instances, an epileptic seizure. However, because epilepsy and tic disorders may develop concomitantly [45-52] and respond to the same treatment [47, 52], the potential presence of epilepsy in our rats does not negate the argument that they also had motor tics. As to whether hyperkinesia was suppressible remained undetermined in our study. However, the fact that hyperkinesia in our rat model was previously shown to subside during sleep suggests that it can be consciously suppressed [24]. Also, another study recently demonstrated that hyperkinesia in our rat model is associated with a decrease in striatal dopamine release [53]. Given the key role striatal dopamine plays in mediating positive and negative reinforcement [54], it is plausible that hyperkinesia in our rats was reinforced by an urge to gain reward and avoid punishment, as often reported by TS patients [55].

As for the nature of vocalizations, their spectral frequency and shape suggest that they were unlikely to reflect the rat’s actual mood, nor were they intended to convey social cues, which is consistent with the notion that phonic tics in TS patients lack meaning [55]. Furthermore, the fact that vocalizations in our rats resulted from disinhibition of NAc, but not CPu, aligns with clinical observations that normal speech is associated with dopamine release in CPu [56], while stuttering is associated with structural alterations in NAc [57].

We also show that DBS of CM/VL attenuates hyperkinesia. Given that rat CM/VL approximates clinical DBS targets [8], our study further validates striatal disinhibition as a TS model. Although the evidence for this comes only from rats that received bicuculline in CPu, hyperkinesia in the NAc group most likely resulted from the spread of bicuculline towards CPu, as discussed above. Therefore, it is likely that DBS of CM/VL could also attenuate hyperkinesia in the NAc group if it were to develop consistently with each bicuculline injection, allowing for the assessment of the DBS effect with greater statistical power. Interestingly, in the CPu group, DBS of CM/VL showed stronger attenuation of hyperkinesia when tested at a lower intensity. Given that a larger intensity increases the spread of the electric field [32] and the number of activated neurons [58], this indicates that a lower intensity was more efficacious because it was more selective in targeting CM/VL. This notion is supported by our finding that DBS of the adjacent regions failed to attenuate hyperkinesia in the CPu group.

Additionally, our study provides early evidence that DBS of CM/VL attenuates vocalizations resulting from NAc disinhibition. However, in contrast to its effect on hyperkinesia, the effect on vocalizations was stronger at a higher intensity. This indicates that the mechanisms for attenuation of hyperkinesia and vocalizations differ, and that while CM/VL may represent the optimal target for attenuation of hyperkinesia, DBS of the adjacent regions may be more suitable for attenuation of vocalizations. This aligns with clinical findings that DBS of the same thalamic region may attenuate motor and phonic tics with different efficacy [59, 60] due to activation of different brain networks [61].

Although the exact mechanisms remained uncertain, DBS appeared to directly oppose bicuculline action rather than induce compensatory responses. This assumption stems from our observation that DBS did not alter arrhythmicity nor the pattern in which hyperkinesia developed. Moreover, it did not affect the normal behavior of rats that received aCSF. Instead, DBS appeared to first fragment dystonia into myoclonia and then attenuate the latter. Also, it has recently been shown that hyperkinesia in our rat model can be attenuated by DBS of the parafascicular nucleus (Pf), which acts by promoting striatal dopamine release [53]. While DBS of Pf and DBS of CM/VL innervate adjacent regions, the degree of similarity between their underlying mechanisms remains to be determined [8].

In conclusion, our study confirms previous findings that bicuculline-induced striatal disinhibition induces hyperkinesia in rats. Furthermore, while previously shown only in monkeys, we demonstrate that it also induces vocalizations in rats. Given the similarity of hyperkinesia to motor tics and vocalizations to phonic tics, we argue that bicuculline-induced striatal disinhibition can serve as a TS rat model. The model’s responsiveness to DBS makes it particularly valuable for unraveling the mechanism of action behind existing DBS protocols as well as for facilitating the development of new ones for symptom-based TS therapy.

## Acknowledgements

This work was supported by the Faculty of Medicine of the University of Cologne (Köln Fortune Programme, 327/2018). VVV, JCB, and TSc are funded by the German Research Foundation (*Deutsche Forschungsgemeinschaft (DFG)*, SFB-1451).

## Legends

Video S1. *Example of vocalizations in conjunction with hyperkinesia*. The audio signal was high-pass filtered at 20 kHz to remove noise from paws scratching the floor. Ultrasound frequencies were then downsampled to the human hearing range. The video file is available on the bioRxiv website.

